## Supplemental material for "Domestic dogs maintain positive clinical, nutritional, and hematological health outcomes when fed a commercial plant-based diet for a year"

### SUPPLEMENTARY MATERIAL

**SUPPLEMENTARY TABLE 1**. Complete blood count and blood chemistry analyses in dogs consuming meat-based diets (baseline) versus plant-based nutrition (6 and 12 months). Values reflect median (minimum - maximum).

CBC values remained within normal reference intervals. In one dog, the total leukocyte count was within the normal interval, but the neutrophil count (2.5 K/μL) was below the lower end of the reference interval at 6 months, which normalized at 12 months. Standard blood chemistry values all remained within clinically unremarkable reference intervals. One dog had a borderline low phosphorous level (2.2 mg/dL) at the endpoint (12 months) in combination with a normal vitamin D level (237 nmol/L). The sodium level (130 mmol/L) measured below the minimum value of the reference interval at baseline in one dog, which normalized at 6 and 12 months. The chloride level was borderline low (108 mmol/L) in one dog at the endpoint. The albumin level (4.3 g/dL) measured above the maximum value of the reference interval in one dog at endpoint, while total protein remained within the normal reference interval. GGT (gamma-glutamyl transferase) baseline levels were above the maximum value of the reference interval in two dogs (13 and 19 U/L), which had normalized at 6 and 12 months. The bilirubin level (1.2 mg/dL) was mildly elevated at baseline in one dog, which normalized at 6 and 12 months. Osmolality measured below (268 mmol/kg) and above (313 mmol/kg) the reference interval in two dogs at baseline, but normalized at 6 and 12 months.

| Parameter | Unit | Baseline | 6 months | 12 months | P-value (Friedman) | P-value (Wilcoxon) | Reference interval (IDEXX) |
|---|---|---|---|---|---|---|---|
| *CBC* | | | | | | | |
| Erythrocytes | M/µL | 7.6 (6.8-8.5) | 7.9 (6.8-8.7) | 8.0 (7.4-8.9) | 0.03 | 0.003 | 5.65-8.87 |
| Hematocrit | % | 52.0 (46.0-58.1) | 53.1 (47.5-58.4) | 54.1 (48.1-60.7) | 0.002 | 0.007 | 37.3-61.7 |
| Hemoglobin | g/dL | 19.1 (16.6-20.5) | 19.2 (17.3-20.7) | 19.6 (17.9-21.2) | 0.003 | 0.002 | 13.1-20.5 |
| Leukocytes | K/µL | 7.3 (5.2-11.9) | 7.6 (4.8-13.3) | 7.0 (5.0-12.1) | 0.59 | 0.73 | 5.05-16.76 |
| Neutrophils | K/µL | 4.7 (3.6-9.1) | 5.2 (2.5-10.5) | 4.6 (3.4-8.8) | 0.89 | 0.56 | 2.95-11.64 |
| Lymphocytes | K/µL | 1.7 (0.9-2.2) | 1.8 (1.2-3.1) | 1.7 (1.0-2.1) | 0.30 | 0.22 | 1.05-5.10 |
| Monocytes | K/µL | 0.4 (0.2-0.9) | 0.4 (0.2-0.8) | 0.4 (0.3-0.9) | 0.69 | 0.24 | 0.16-1.12 |
| Eosinophils | K/µL | 0.4 (0.3-1.0) | 0.4 (0.2-1.0) | 0.5 (0.3-1.2) | 0.62 | 0.35 | 0.06-1.23 |
| Basophils | K/µL | 0.01 (0.00-0.09) | 0.01 (0.00-0.08) | 0.02 (0.00-0.06) | 0.69 | > 0.99 | 0.00-0.10 |
| Platelets | K/µL | 202 (144-322) | 232 (140-400) | 214 (185-318) | 0.34 | 0.21 | 148-484 |
| *Chemistry* | | | | | | | |
| Glucose | mg/dL | 99 (76-109) | 99 (75-109) | 107 (85-117) | 0.01 | < 0.001 | 74-143 |
| Creatinine | mg/dL | 1.0 (0.6-1.4) | 1.1 (0.8-1.4) | 1.1 (0.7-1.3) | 0.14 | 0.13 | 0.5-1.8 |
| BUN | mg/dL | 20 (9-23) | 15 (6-23) | 19 (6-27) | 0.09 | 0.77 | 7-27 |

| | | | | | | | |
|---|---|---|---|---|---|---|---|
| Phosphorus | mg/dL | 4.4 (3.2-5.8) | 4.0 (2.5-6.0) | 3.9 (2.2-4.8) | 0.08 | 0.01 | 2.5-6.8 |
| Calcium | mg/dL | 10.2 (9.8-10.6) | 10.1 (9.6-10.6) | 10.2 (9.1-10.5) | 0.21 | 0.36 | 7.9-12.0 |
| Sodium | mmol/L | 153 (130-158) | 153 (146-156) | 150 (145-152) | 0.002 | 0.01 | 144-160 |
| Potassium | mmol/L | 4.2 (3.6-5.0) | 4.4 (3.8-5.5) | 4.6 (4.2-5.5) | < 0.001 | 0.001 | 3.5-5.8 |
| Chloride | mmol/L | 117 (112-125) | 115 (110-119) | 113 (108-116) | 0.03 | 0.004 | 109-122 |
| Total Protein | g/dL | 6.7 (6.0-7.4) | 6.9 (6.1-7.7) | 6.9 (6.1-7.4) | 0.27 | 0.97 | 5.2-8.2 |
| Albumin | g/dL | 3.1 (2.7-3.5) | 3.4 (3.0-3.9) | 3.6 (3.1-4.3) | 0.002 | 0.003 | 2.3-4.0 |
| ALT | U/L | 97 (30-129) | 61 (22-115) | 51 (21-357) | 0.06 | 0.12 | 10-125 |
| ALP | U/L | 61 (13-222) | 44 (22-222) | 40 (10-192) | 0.15 | 0.05 | 23-212 |
| GGT | U/L | 3 (0-19) | 1 (0-7) | 3 (0-6) | 0.003 | 0.62 | 0-11 |
| Bilirubin | mg/dL | 0.4 (0.1-1.2) | 0.4 (0.2-0.6) | 0.2 (0.1-0.5) | 0.02 | 0.006 | 0.0-0.9 |
| Cholesterol | mg/dL | 178 (121-290) | 210 (111-288) | 193 (131-280) | 0.52 | 0.40 | 110-320 |
| Osmolality | mmol/kg | 303 (268-313) | 303 (289-310) | 301 (291-305) | 0.24 | 0.12 | 290-310 |
| T4 | µg/dL | 1.7 (0.9-2.6) | 1.5 (1.0-3.7) | 1.8 (1.1-3.7) | 0.42 | 0.32 | 1.0-4.0 |

**SUPPLEMENTARY TABLE 2.** Urinalysis in dogs consuming meat-based diets (baseline) versus plant-based nutrition (6 and 12 months). Values refer to median (minimum - maximum). N/A: not applicable (since each row has zero difference, which precludes calculation of a paired test). Semi-quantitative UA parameters were tabulated as ‘0’ (normal/negative/not detected or <1/HPF), ‘1’ (trace), or the highest reported value (e.g., 1-5/HPF and 6-20/HPF were tabulated as ‘5’ and ‘20’, respectively), where HPF is high power field. Urine samples were collected via cystocentesis where iatrogenic microscopic hematuria is to be expected.

Urine pH trended downwards and towards normal - although findings did not reach statistical significance (p = 0.05) - as levels were elevated in eight dogs at baseline (pH 8-9), four dogs at 6 months (pH 8) and two dogs at endpoint (pH 8). Potential crystal formation is associated with changes in urine pH. In this study, we identified a variety of crystals in the urine from a total of 60% of the dogs (9 of 15) at different time points (including 3 dogs at baseline, 3 different dogs at 6 months, and 3 different dogs at 12 months) with no identifiable pattern to the changes.

| Parameter | Baseline | 6 months | 12 months | P-value (Friedman) | P-value (Wilcoxon) | Normal Values |
|---|---|---|---|---|---|---|
| Specific gravity | 1.044 (1.020-1.050) | 1.036 (1.015-1.050) | 1.036 (1.015-1.050) | 0.06 | 0.06 | Variable |
| Glucose | 0 (0-0) | 0 (0-100) | 0 (0-0) | 0.37 | N/A | Negative |
| Bilirubin | 0 (0-1) | 0 (0-1) | 0 (0-1) | 0.37 | 0.63 | Negative |
| Ketones | 0 (0-15) | 0 (0-15) | 0 (0-15) | 0.20 | N/A | Negative |
| pH | 8 (5-9) | 7 (5-9) | 7 (5-8) | 0.05 | 0.03 | 5.0-7.5 |
| Protein | 1 (0-30) | 1 (0-30) | 1 (0-30) | 0.61 | 0.31 | Negative |
| Heme | 0 (0-50) | 10 (0-250) | 0 (0-25) | 0.01 | 0.14 | Negative |
| Erythrocytes | 0 (0-14) | 1 (0-50) | 0 (0-4) | 0.66 | 0.26 | 0-5/HPF |
| Leukocytes | 0 (0-17) | 0 (0-9) | 0 (0-3) | 0.63 | 0.44 | 0-5/HPF |
| Casts | 0 (0-0) | 0 (0-1) | 0 (0-0) | 0.14 | N/A | None |
| Epithelial cells | 0 (0-2) | 0 (0-5) | 0 (0-2) | 0.20 | 0.63 | Variable |
| Crystals | 0 (0-20) | 0 (0-5) | 0 (0-50) | 0.72 | 0.44 | None |
| Bacteria | 0 (0-1) | 0 (0-0) | 0 (0-0) | 0.37 | > 0.99 | None |

**SUPPLEMENTARY TABLE 3.** Nutrient analysis including plasma amino acid and serum L-carnitine concentrations in dogs consuming meat-based diets (baseline) versus plant-based nutrition (6 and 12 months). Values refer to median (first quartile - third quartile). Reference values for first and third quartiles were provided by UCD. We requested reference intervals (minimum to maximum) from this laboratory, but were informed they were not available.

When evaluating the entire data set (as shown in Figure 2), some data points were below the $Q_1$ for six of the ten *essential* AAs. We initially extrapolated the reference interval (minimum to maximum) for the essential amino acid tryptophan (11 – 103 nmol/L) based on the first ($Q_1$) and third ($Q_3$) quartiles, and the interquartile range (IQR), as follows. If $IQR = Q_3 - Q_1$, the maximum can be calculated as ($Q_3 + 1.5 \times IQR = 68 + 1.5 \times 23 = 103$) and the minimum as ($Q_1 – 1.5 \times IQR = 45 – 1.5 \times 23 = 11$). We moreover used extrapolation to calculate minimum values for arginine (28), leucine (36.5), methionine (15), threonine (28.5), and valine (56.5), as shown in Figure 2.

Three of the *essential* AA levels were significantly different between baseline and endpoint, including methionine ($p = 0.02$), phenylalanine ($p = 0.01$), and tryptophan ($p = 0.01$). Values were either within or above the interquartile reference intervals (first quartile - third quartile) provided by UCD, and above the minimum values derived through extrapolation. The values for all three of these AAs trended upwards with higher values at endpoint compared to baseline.

Statistically significant differences were found in the levels of five *non-essential* AAs between baseline and endpoint, including alanine ($p < 0.001$), cystathionine ($p < 0.001$), glutamate ($p = 0.02$), glutamine ($p < 0.001$), serine ($p = 0.03$), and tyrosine ($p = 0.001$). All measured within or above the reference intervals and values for all five AAs exhibited an upwards trend. Another four *non-essential* AAs (cysteine, hydroxyproline, ornithine, proline) measured below the first quartile, but showed no statistically significant differences between baseline and endpoint values. Cysteine levels were low, or below detection, in most samples (which is commonly attributed to storage loss). Hydroxyproline and proline levels increased over time, while ornithine levels decreased slightly.

| **Amino acid** | **Baseline (nmol/ml)** | **6 months (nmol/ml)** | **12 months (nmol/ml)** | **P-value (Friedman)** | **P-value (Wilcoxon)** | **Ref. Values (UC Davis)** |
|---|---|---|---|---|---|---|
| *Essential* | | | | | | |
| Arginine | 118 (112-144) | 103 (99-109) | 127 (112-164) | < 0.001 | 0.72 | 85-123 |
| Histidine | 84 (79-89) | 86 (84-88) | 94 (88-105) | 0.01 | 0.07 | 60-80 |
| Isoleucine | 84 (73-90) | 64 (58-68) | 75 (61-82) | 0.001 | 0.22 | 40-57 |
| Leucine | 134 (114-154) | 125 (111-129) | 138 (127-179) | 0.03 | 0.12 | 95-134 |
| Lysine | 149 (140-160) | 156 (140-169) | 167 (148-238) | 0.13 | 0.09 | 94-159 |
| Methionine | 51 (48-56) | 51 (49-53) | 68 (51-78) | 0.01 | 0.02 | 45-65 |
| Phenylalanine | 61 (59-64) | 72 (66-73) | 72 (66-79) | 0.002 | 0.01 | 39-52 |
| Threonine | 172 (136-298) | 162 (151-185) | 226 (207-246) | 0.03 | 0.60 | 138-211 |
| Tryptophan | 36 (24-42) | 35 (34-42) | 52 (39-75) | 0.01 | 0.01 | 45-68 |
| Valine | 173 (137-201) | 155 (145-158) | 179 (157-214) | 0.04 | 0.32 | 130-179 |
| *Non-essential* | | | | | | |
| Alanine | 396 (352-432) | 562 (551-588) | 578 (533-651) | < 0.001 | < 0.001 | 320-455 |
| Asparagine | 63 (50-73) | 45 (42-49) | 55 (47-96) | 0.01 | 0.95 | 30-49 |
| Aspartate | 11 (9-13) | 9 (6-9) | 10 (9-15) | 0.02 | 0.79 | 6-8 |
| Butyrate | 22 (18-24) | 32 (25-34) | 17 (15-27) | 0.06 | 0.57 | ND |
| Citrulline | 52 (45-56) | 36 (35-44) | 53 (34-65) | 0.04 | 0.92 | 27-50 |
| Cystathionine | 5 (5-6) | 8 (7-9) | 8 (8-10) | < 0.001 | < 0.001 | ND |
| Cysteine | 0 (0-1) | 0 (0-4) | 1 (0-1) | 0.32 | 0.34 | 36-53 |

| | | | | | | |
|---|---|---|---|---|---|---|
| Glutamate | 72 (59-84) | 97 (74-124) | 91 (75-106) | 0.01 | 0.02 | 15-26 |
| Glutamine | 564 (539-590) | 643 (626-683) | 735 (693-773) | < 0.001 | < 0.001 | 417-569 |
| Glycine | 201 (185-243) | 185 (174-187) | 241 (195-351) | 0.01 | 0.38 | 207-310 |
| Hydroxyproline | 21 (16-33) | 12 (9-18) | 29 (18-51) | 0.01 | 0.22 | 44-78 |
| Methylhistidine1 | 11 (9-14) | 6 (4-9) | 8 (3-13) | 0.01 | 0.08 | ND |
| Methylhistidine3 | 5 (4-7) | 2 (1-4) | 5 (3-6) | 0.01 | 0.45 | ND |
| Ornithine | 19 (9-27) | 14 (12-16) | 15 (13-18) | 0.62 | 0.19 | 23-43 |
| Proline | 161 (112-201) | 147 (129-172) | 202 (161-206) | 0.08 | 0.15 | 174-304 |
| Serine | 121 (114-128) | 121 (112-125) | 137 (122-161) | 0.01 | 0.03 | 87-126 |
| Taurine | 106 (79-132) | 89 (78-92) | 136 (115-152) | < 0.001 | 0.06 | 60-90 |
| Tyrosine | 34 (31-39) | 44 (39-52) | 53 (47-58) | < 0.001 | 0.001 | 30-47 |
| L-Carnitine | 19 (13-21) | 27 (17-31) | 35 (14-43) | 0.09 | 0.06 | ND |

**SUPPLEMENTARY TABLE 4.** Nutrient analysis of serum vitamin concentrations in dogs consuming meat-based diets (baseline) versus plant-based nutrition (6 and 12 months). Values refer to median (minimum - maximum).

| Vitamin | Unit | Baseline | 6 months | 12 months | P-value (Friedman) | P-value (Wilcoxon) | Ref. Values (MSU/TAMU) |
|---|---|---|---|---|---|---|---|
| *Lipid-soluble* | | | | | | | |
| Vitamin A | ng/mL | 734 (437-1,295) | 924 (600-1,292) | 979 (769-1,330) | 0.01 | 0.01 | 400-1,200 |
| Vitamin D | nmol/L | 120 (57-418) | 228 (91-383) | 257 (173-418) | < 0.001 | 0.004 | 109-423 |
| Vitamin E | ug/mL | 42 (21-74) | 31 (25-56) | 37 (25-57) | 0.62 | 0.33 | 4-12 |
| *Water-soluble* | | | | | | | |
| Folate (B9) | ug/L | 9 (5-23) | 11 (4 -15) | 11 (4-37) | 0.04 | 0.09 | 8 – 24 |
| Cobalamin (B12) | ng/L | 364 (261-639) | 427 (310-619) | 426 (278-505) | 0.25 | 0.17 | 251 - 908 |

**ADDITIONAL ACKNOWLEDGEMENTS**

The authors would like to thank the organizing team behind the Plant-Based Dog Food Health Study Initiative in Los Angeles, California for facilitating the fundraising efforts that made this study a reality. We thank the foundations and individual donors all of whom contributed generously. We acknowledge research support from the WesternU College of Veterinary Medicine Office for Research, specifically Christiana Benoit, Trinidad Cisneros, Dominique Griffon, and the True One Medicine Initiative. The authors are grateful to colleagues and students at the WesternU PHC who provided technical support during a long-term study conducted under pandemic restrictions, especially Brittany Bryan, Annette Chavarria-Marron, Suzanna Corral, Jennie Jennings, Zach Morris, and Andres Muñoz. We also acknowledge BioNote, Inc. for providing the Vcheck200 Analyzer and the canine NT-proBNP assays for this study. The authors wish to acknowledge T. Colin Campbell of Cornell University, Hana Khaleova of the Physicians Committee for Responsible Medicine, and Denis M. Medeiros of the University of Missouri for sharing their wealth of nutritional science expertise and clinical study design experience. Our team is sincerely grateful to the dogs who were enrolled in this clinical study and the clients who enabled their participation despite challenging restrictions imposed by the Covid19 pandemic.
