## Supplemental Figure 1 for "Domestic dogs maintain positive clinical, nutritional, and hematological health outcomes when fed a commercial plant-based diet for a year"

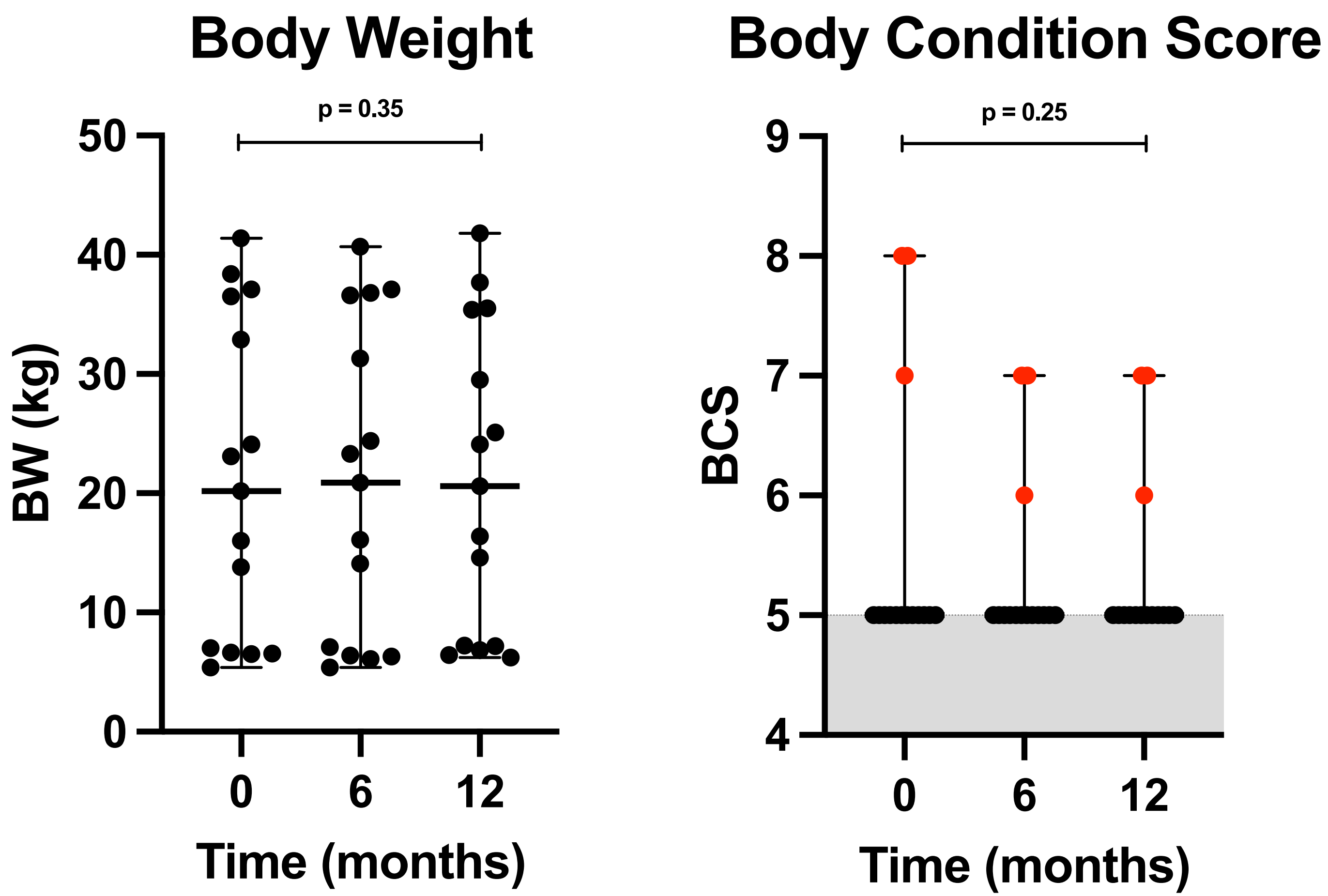


**Supplemental Figure 1.** Scatter plots of body weight and body condition score (BCS) in dogs at 0, 6, and 12 months. Black and red data points represent dogs with normal and increased BCS, respectively. The grey-shaded area represents normal range for BCS (4-5).
